# SFRP1 drives glycolytic activation in astrocytes during neuroinflammation

**DOI:** 10.1101/2025.09.10.675368

**Authors:** Marcos Martínez-Baños, Pablo Miaja Hernández, María Jesús Martin-Bermejo, Guadalupe Pereyra, Paola Bovolenta

**Affiliations:** Centro de Biología Molecular Severo Ochoa, CSIC-UAM, Madrid, Spain

**Keywords:** astrocytes, inflammation, metabolism, microglia, mitochondria

## Abstract

Astrocytes and microglia maintain brain homeostasis and respond to inflammation through functions coordinated by molecular mediators they produce. Growing evidence shows that cellular metabolism is key to how these cells adapt to challenges. However, little is known about what drives glial metabolic reprogramming or whether molecules involved in astrocyte– microglia crosstalk also regulate this process. Here, we explored this question focusing on Secreted Frizzled-Related Protein 1 (SFRP1). SFRP1 is an astrocyte-derived factor induced by inflammatory cues and overexpressed in neurodegeneration, which fosters microglial response to inflammation through NF-κB/HIF-dependent programs. We combined mitochondrial morphometry (MitoTracker Red and MiNA analysis) with Seahorse extracellular flux assays (Mito Stress Test) to determine whether SFRP1 modulates glial bioenergetics in primary cultures of astrocytes and microglia from wild-type and *Sfrp1*^*-/-*^ mice. We report that SFRP1 acts as a driver of astrocytic metabolic activation, preferentially enhancing glycolysis over mitochondrial respiration. This effect is most pronounced during inflammation, when oxidative phosphorylation is restricted and SFRP1 enhances glycolytic flexibility to sustain energy demands. By contrast, microglia showed the expected LPS-driven glycolytic shift with minimal dependence on SFRP1 under monoculture conditions. These findings position SFRP1 as a candidate regulator of astrocyte-centered metabolic tuning during neuroinflammation, with implications for disorders such as Alzheimer’s disease, in which SFRP1 is elevated.

## Introduction

Astrocytes, the most abundant glial cell type, and microglia, the resident immune cells of the brain, act as key regulators of Central Nervous System (CNS) homeostasis. Both cell types participate in diverse processes, including synaptic modulation, clearance of cellular debris, maintenance of blood-brain barrier integrity, and provide metabolic and functional support to the neurons. They also play a critical role in the rapid initiation of inflammatory response upon CNS damage or infection (Singh, 2022). Growing evidence indicates that cellular metabolism is a major determinant of the capacity of both astrocytes and microglial cells to adapt to environmental challenges, thereby shaping their functional states under both physiological and pathological conditions (Afridi et al., 2022). Metabolic reprogramming in these cells is particularly critical during neuroinflammation, as it influences the production of pro- and anti-inflammatory mediators, which, in neurodegenerative diseases, ultimately modulate disease progression (Afridi et al., 2022).

In their patrolling state, microglia primarily rely on oxidative phosphorylation and glucose oxidation to generate ATP via mitochondrial respiration (Bernier, York, Kamyabi, et al., 2020). This metabolic profile supports their energy-demand in a low-inflammatory status, while also regulating redox balance and the production of immunomodulatory molecules (Bernier, York, Kamyabi, et al., 2020). Upon activation, however, microglial cells shift their metabolism toward aerobic glycolysis, enabling faster biosynthetic intermediates production to sustain active phagocytosis, cytokine release and proliferation. Although this metabolic reprogramming is essential for mounting a fast immune response, chronic activation locks microglial cells in a glycolytic, pro-inflammatory state, which fosters a neurotoxic environment marked by aberrant release of proinflammatory cytokines (TNF-α, IL-1β, IL-6), complement proteins (Hoogland et al., 2015; Hong et al., 2016) and reactive oxygen species (ROS), ultimately driving synaptic degradation and neuronal damage.

Astrocytes, by contrast, rely predominantly on aerobic glycolysis, providing metabolic support to neurons through lactate production via the astrocyte-neuron lactate shuttle. This metabolic coupling ensures a continuous energy supply for neuronal activity (Barros & Weber, 2018). Astrocytes also safeguard the brain against oxidative stress by synthesizing and releasing glutathione precursors and serve as a local energy reservoir owing to their glycogen storage capacity, which can be mobilized in response to increased energy demands or metabolic stress (Bak et al., 2018). Under inflammatory conditions, such as those associated with neurodegenerative diseases or LPS stimulation, astrocytes reduce their glycolytic activity (Robb et al., 2020), develop mitochondrial dysfunction and exhibit impaired lactate export, collectively compromising neuronal support. Impairment of glutamate uptake and increased oxidative stress, exacerbates neuronal vulnerability and synaptic dysfunction (Bak et al., 2018), while reactive astrocytes further amplify neuroinflammation by releasing pro-inflammatory mediators that stimulate the activation of neighboring microglia (Liddelow et al., 2017).

NF-κB and HIF1α signaling drive metabolic adaptations of both astrocyte and microglia (Ishiyama et al., 2025; Madai et al., 2024; Robb et al., 2020). For example, pharmacological inhibition of NF-κB blocks LPS-induced metabolic reprogramming, reducing glycolysis, mitochondrial respiration, and cytokine release (TNF-α, IL-6, IL-10) (Robb et al., 2020). Conversely, HIF1α activation drives astrocytic glycolysis by upregulating GLUT and LDHA and inhibiting pyruvate oxidation via PDK1 (Brix et al., 2012; Madai et al., 2024; Semenza, 1996). Notably, astrocyte and microglial metabolic adaptations are not independent events. Increasing evidence indicates that the well-established astrocyte-microglia crosstalk involves a tightly coupled metabolic dialogue, in which metabolites such as lactate or reactive oxygen species, together with cytokines and extracellular vesicles act as signaling mediators of this interplay (Bernier, York, & MacVicar, 2020; Hasel et al., 2023; Jha et al., 2019). This dialogue is particularly relevant in neurodegenerative diseases, where the identification of specific metabolic mediators may provide novel therapeutic targets (Hasel et al., 2023).

In this context, we have recently demonstrated that Secreted Frizzled-Related Protein 1 (SFRP1) is an astrocyte-derived factor with a strong impact on both microglial and neuronal function (Pereyra et al., 2025; Rueda-Carrasco et al., 2021). Initially identified as a negative modulator of Wnt signaling, SFRP1 can also interact with and modulate the activity of multiple proteins (Bovolenta et al., 2008), including the metalloprotease ADAM10 (Esteve et al., 2011). While SFRP1 is highly expressed in cortical radial glial progenitors during development (Donega et al., 2022; Esteve, Crespo, et al., 2019), in the adult brain its expression remains restricted to the choroid plexus, ventricular neurogenic niches and, at low levels, in astrocytes (Esteve et al., 2019). Under neurodegenerative conditions, however, astrocytes upregulate SFRP1 (Esteve et al., 2019) and the protein markedly accumulates in the brain and CSF of Alzheimer’s disease patients (Askenazi et al., 2023; Esteve, et al., 2019; Higginbotham et al., 2020; Johnson et al., 2022). A similar induction occurs after LPS injection in the mouse brain or following LPS treatment of cultured astrocytes (Rueda-Carrasco et al., 2021). Transcriptomic analysis of microglia isolated from wild-type (WT) and *Sfrp1*^-/-^ mouse after LPS exposure revealed that SFRP1 fosters activation of the hypoxia-inducible factor (HIF) and NF-κB inflammatory pathways in microglial cells (Rueda-Carrasco et al., 2021). Both pathways are central regulators of glial metabolic reprograming and drive the expression of glycolytic and redox genes in response to inflammation (Corcoran & O’Neill, 2016). Consistent with this notion, a number of HIF and NF-κB target genes involved in metabolic processes, including *Ldha, Sod2* and *Aldoa*, were significantly downregulated in SFRP1 absence (Rueda-Carrasco et al., 2021).

These findings suggest a mechanistic link between SFRP1 activity and metabolic reprogramming of glial cells during inflammation. To explore this possibility, we performed Seahorse-based metabolic profiling of cultured astrocytic and microglial cells and found that SFRP1 contributes to metabolic reprogramming of activated astrocytes by selectively enhancing glycolytic activity and promoting their entrance into an energetic state.

## Materials and Methods

### Mice

C57BL/6J wild type and *Sfrp1*^-/-^ mice were used to generate astrocytes and microglial cultures. *Sfrp1*^-/-^ mice were obtained by outcrossing *Sfrp1*^-/-^;*Sfrp2*^+/-^ (Satoh et al., 2006) in a 129/C57BL/6 background, followed by back-crossing with C57BL/6J to clean the background. Mice were housed in the animal facility of the Centro de Biología Molecular Severo Ochoa in a temperature and humidity-controlled, pathogen-free environment, with 12-hour light/dark cycles and *ad libitum* feeding-drinking. All animal procedures were approved by the ethical committees of the CSIC and Comunidad Autónoma de Madrid.

### Primary mixed glial cultures

Primary glial cultures were established using cerebral cortices of postnatal P2-P4 WT and or *Sfrp1*^-/-^ mice. Cortices were dissected in Ca^2+^ and Mg^2+^ free Hank’s Balanced Salt Solution (HBSS; Invitrogen). Tissue was chopped and mechanically dissociated with a Pasteur pipette, treated with DNase I (50 µg/ml; Sigma-Aldrich, Cat. No. DN25) in HBSS for 10 min at 37ºC. Cell suspension was centrifuged at 200 g for 8 min. The pellet was resuspended in Dulbecco’s modified Eagle medium and F-12 nutrient mixture (DMEM/F12; Invitrogen) containing 10% Fetal Bovine Serum (FBS; Invitrogen), and penicillin/streptomycin (Sigma Aldrich). Cortices from 3 pups were platted in 75 cm^2^ flaks, pre-treated with Poly-D-Lys (10 μg/mL, Sigma Aldrich, Cat. No. P7280) in 0.1M borate buffer. Cells were cultured at 37ºC in a humidified 5% CO2 incubator. After two days, the medium was replaced with fresh medium supplemented with 10% conditioned medium from the m-CSF1-producing L929 cell line to enhance microglial survival. Mixed glial cultures were maintained until confluence, which was typically reached within 2 weeks without medium changes. Astrocytes and microglia were separated from the mixed cultures by mechanical agitation (shaking and gentle beating for 15 min), which induced the selective detachment and floating of microglial cells into the culture medium. After microglia collection, culture flasks were washed with PBS and incubated with 0.25% trypsin (Gibco, Life Technologies, Cat. No. 25300054), 1mM EDTA in PBS for 5min at 37ºC to detach adherent astrocytes. Trypsin activity was halted by the addition of DMEM with 10% FBS. Astrocyte and microglia cell suspensions were centrifuged at 250 g for 8 min. Microglial supernatant was stored for further use. The cell pellets were resuspended in DMEM/F12 (500 µL) and viable cells were counted. Cells were seeded into appropriate culture plates at the desired density in 50% fresh DMEM/F12 and 50% of the recovered medium.

### Seahorse

Mitochondrial respiration was assessed using the Seahorse XF96 Extracellular Flux Analyzer (Agilent Technologies), which measures oxygen consumption (OCR) and extracellular acidification (ECAR) rates of live cells (Mookerjee et al., 2015). Purified astrocyte and microglial cells from WT or *Sfrp1*^-/-^ were seeded at a density of 80,000 and 40,000 cells per well, respectively, in Seahorse XF96 cell culture microplates (Agilent, Cat. No. 103792-100) pre-treated with Poly-D-Lys. After two days of incubation, cells were treated for 24 h with LPS (0.1 µg/ml) and hrSFRP1 protein (0.2 and 0.5 µg/ml). The assay was performed in DMEM without sodium bicarbonate, to ensure optimal pH stability, supplemented with glucose (5 mM; Gibco, Cat. No. A24940-01), sodium pyruvate (1 mM; Invitrogen) and glutamine (2 mM; Invitrogen). After establishing basal respiration rate, the following mitochondrial modulators were sequentially added to evaluate different mitochondrial parameters: oligomycin (1 µM; Sigma-Aldrich, Cat. No. O4876), an inhibitor of ATP synthase (Complex V), to assess ATP-linked respiration; FCCP (Carbonyl cyanide 4-(trifluoromethoxy)phenylhydrazone, 1.5 µM; Sigma-Aldrich, Cat. No. C2920), a mitochondrial uncoupler that collapses the proton gradient and reveals the maximal respiratory capacity under conditions of increased energy demand; a mixture of Rotenone (2 µM; Sigma-Aldrich, Cat. No. R8875) an inhibitor of mitochondrial complex I that impairs ATP production, and antimycin A (2 µM; Sigma-Aldrich, Cat. No. A8674), a complex III blocker that blocks mitochondrial respiration, to measure non-mitochondrial oxygen consumption. Key parameters of mitochondrial function, including basal respiration, maximal respiratory capacity, spare respiratory capacity, proton leak, and ATP-linked respiration, were calculated from OCR values obtained throughout the assay.

### Mitochondrial network analysis

Mitochondria were labelled using MitoTracker Red (ThermoFisher Scientific, Cat. No. 7513) and analyzed using the “Mitochondrial Network Analysis” (MiNA) plugin for ImageJ (Valente et al., 2017). Primary purified cultures of astrocyte and microglia were seed in µ-Slide 8 Well high plates (Ibidi, Cat. No. 80806) at a density of 15,000 cells/cm^2^. After 48 hours of incubation, cells were treated with hrSFRP1 (0.5 ug/mL) for 24 hours. Cells were incubated with Red MitoTracker (2 µM) for 30 min at 37°C and thereafter images were acquired from randomly selected fields using a Zeiss LSM800 vertical laser scanning confocal microscope (Zeiss). Mitochondrial network morphology was evaluated in a minimum of 32 individual cells per experimental condition and six biological replicates using the MiNA plugin.

### Statistical analysis

Statistical analyses were conducted using GraphPad Prism version 9 (GraphPad Software). The statistical tests used and the corresponding p-values are indicated in the figure legends. P-value of <0.05 was considered statistically significant.

## Results

### SFRP1 does not alter mitochondrial architecture of cultured glial cells

The mitochondrial network of individual cells constantly remodels through organelle fission and fusion, a process known as mitochondrial dynamics, which reflects cellular metabolic adaptation to stress conditions (Giacomello et al., 2020; Jackson & Robinson, 2018). Thus, changes in mitochondrial network may reflect alterations in cellular metabolic state. We thus begun by asking if SFRP1 has any influence on the morphology and distribution of mitochondria in cultured astrocytes and microglial cells.

We established purified astrocytes and microglia cultures from WT and *Sfrp1*^-/-^ mice and exposed them to vehicle or recombinant human SFRP1 (hrSFRP1; 500 ng/mL) for 24 hrs. Cultures were then stained with MitoTracker Red and their “mitochondrial skeleton” was established using the Mitochondrial Network Analysis v2.0 toolkit (MiNA; Fig. 1; Valente et al., 2017). This skeleton allows to extract detailed morphological data such as the overall mitochondrial footprint (the total area occupied by mitochondria), the number of individual mitochondria, the number of mitochondrial networks and their average branch length. Analysis of the astrocytic “skeleton” confirmed a mitochondrial distribution similar to that previously reported (Fig. 1A; Ayala et al., 2024; Pasqualotto et al., 2024). On average, each astrocyte contained around 170 mitochondria, arranged into about 15 networks, with branches stretching over 2.1 µm. The total mitochondrial area per cell ranged around 700 µm^2^. Astrocytes from *Sfrp1*^-/-^ mice presented very similar values and hrSFRP1 treatment did not modify any of the assessed parameters (Fig. 1B–E), suggesting that SFRP1 has no impact on astrocytic mitochondrial architecture.

**Figure 1.**
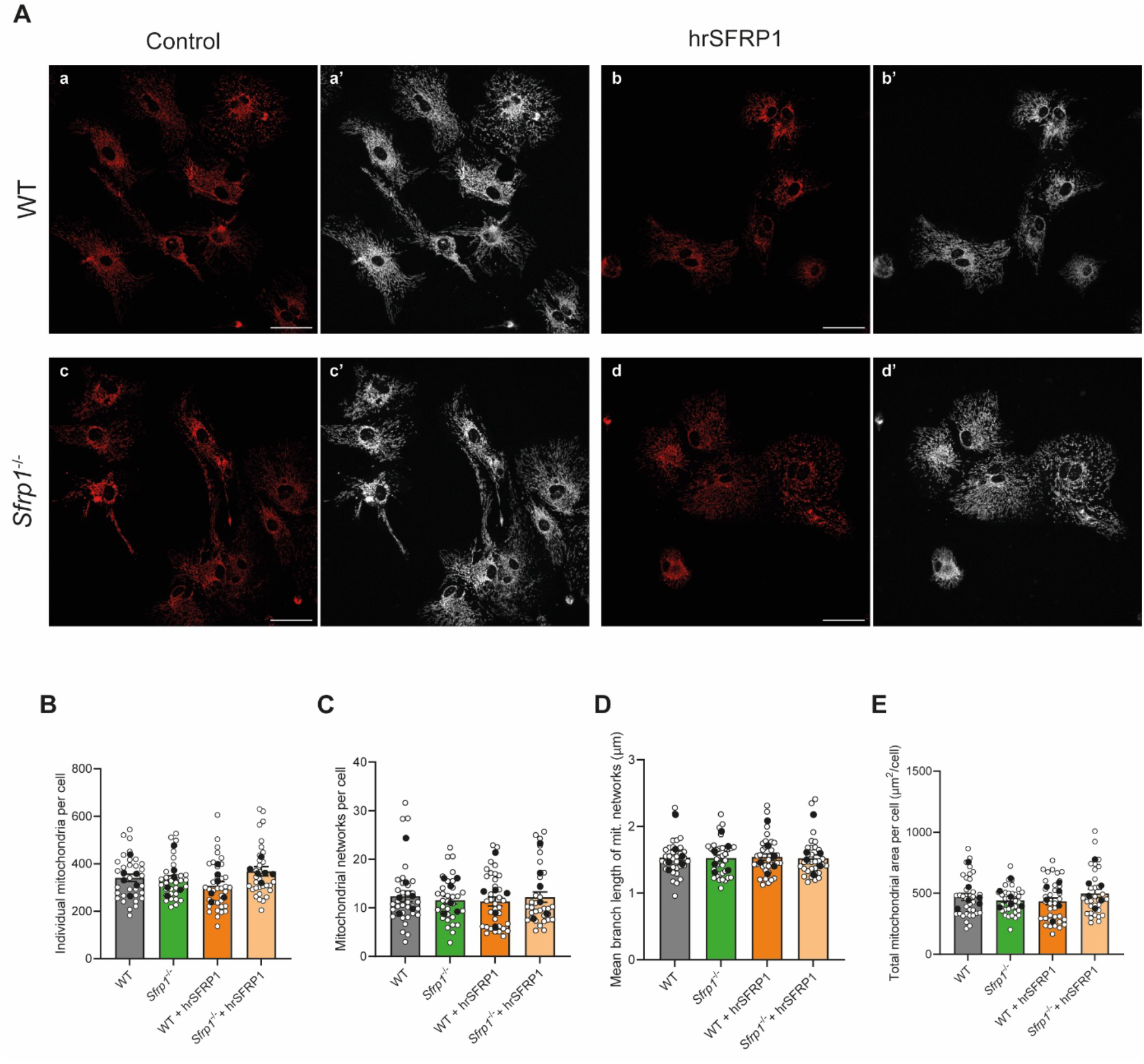
SFRP1 has no influence on astrocytes mitochondrial network. (A) Representative immunofluorescence images of astrocytes, derived from WT and *Sfrp1*^-/-^ mice, treated with or without hrSFRP1 (500 ng/mL) for 24 hr. Cells were stained with Red MitoTracker to visualize mitochondria (a-d). The skeletonized mitochondrial networks (white) was generated with the MiNA toolset for quantitative analysis (a’-d’). Scale bar 50 µm. B-E) The graphs illustrates the number of individual mitochondria per cell (B); mitochondrial network per cell (C); mean branch length of the mitochondrial network (D); and total mitochondrial area per cell (E). Values correspond to mean ± SEM (n = 33-36 acquisitions, white dots; from N = 6 cell cultures, black dots, per group). Statistical comparisons were calculated with one-way ANOVA followed by Tukey multiple comparisons test. No statistical differences were observed.

Similar results were obtained in parallel cultures of microglial cells (Fig. 2), analyzed with the same pipeline. Mitochondrial morphology remained largely unchanged across all experimental conditions (Fig. 2), although microglial cells from *Sfrp1*^-/-^ mice appeared to have slightly fewer individual mitochondria (Fig. 2B), and their network branches were somewhat longer (Fig. 2D). However, addition of hrSFRP1 showed no effect, suggesting that these slight changes may reflect some possible endogenous phenomenon, perhaps only distantly related to SFRP1.

**Figure 2.**
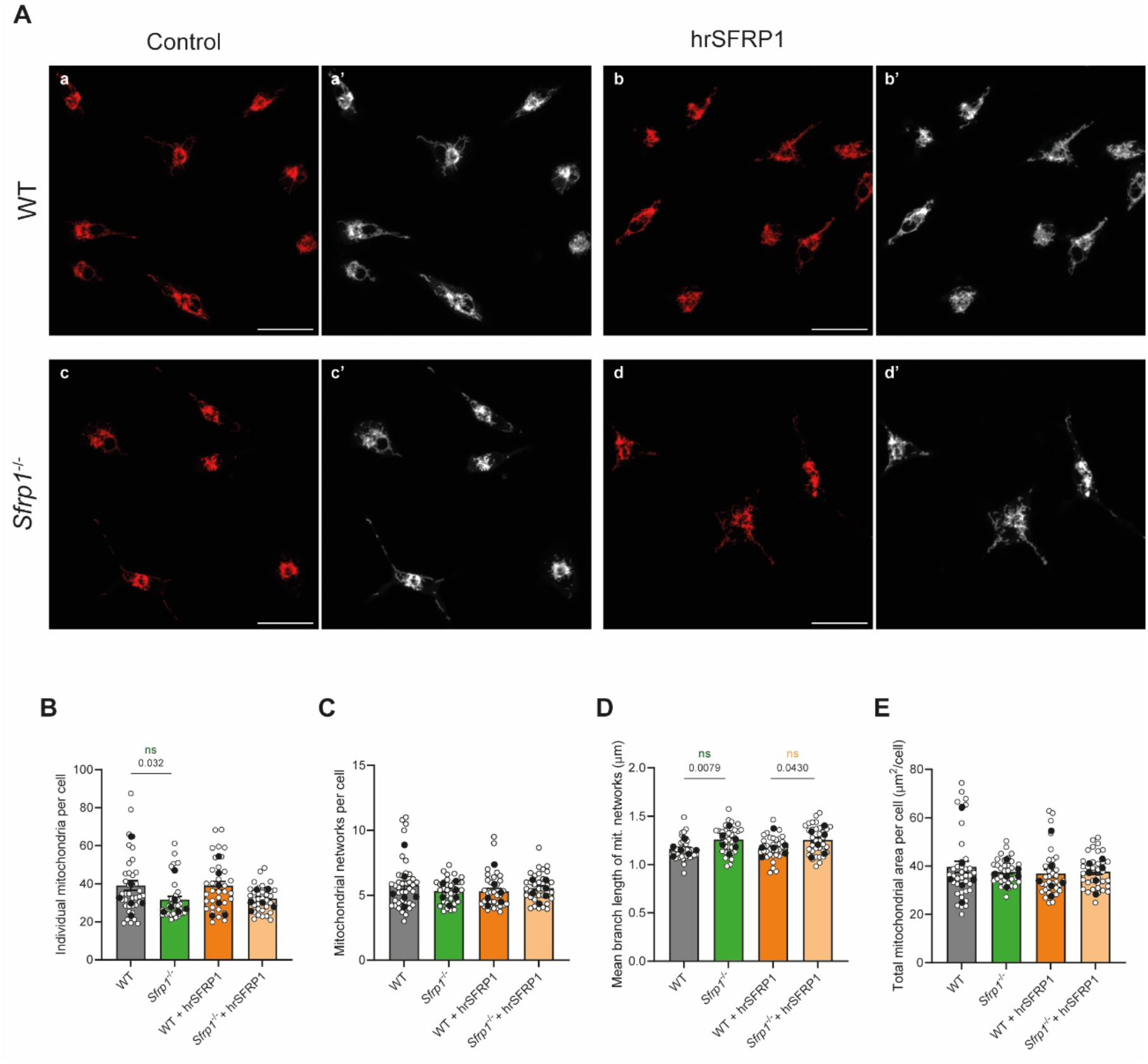
The mitochondrial network is slightly more branched in microglia cells derived from *Sfrp1*^*-/-*^ mice. (A) Representative immunofluorescence images of microglial cultures, derived from WT and *Sfrp1*^-/-^ mice and treated with or without hrSFRP1 (500 ng/mL) for 24 h. Cells were stained with Red MitoTracker to visualize mitochondria (a-d), and skeletonized networks (white) were generated using the MiNA toolset for quantitative analysis (a’-d’). Scale bar 25 µm. Quantification of individual mitochondria per cell (B); mitochondrial network per cell (C); mean branch length of the mitochondrial network (D) and (E) total mitochondrial area per cell. Values in graphs correspond to the mean ± SEM (n = 33-36 acquisitions, white dots; from N = 6 cell cultures, black dots, per group). Statistical comparisons were calculated with one-way ANOVA followed by Tukey multiple comparisons test. WT: Wild-type.

### SFRP1 fosters the shift towards glycolytic metabolism in activated astrocytes

To further analyze the possible impact of SFRP1 on glial metabolism, we turned to Seahorse technology, a powerful tool that tracks mitochondrial performance in real time. This technology measures two key indicators: mitochondrial respiration (oxygen consumption rate, OCR) and glycolytic activity (extracellular acidification rate, ECAR). These parameters are obtained via the “Mito Stress Test,” based on the sequential addition of a series of compounds that interfere with different aspects of mitochondrial function (Fig. 3). These include oligomycin that inhibits ATP production; FCCP, which uncouples the mitochondrial membrane potential, driving cells to their maximal respiration capacity; and finally, rotenone and antimycin A that fully inhibit the electron transport chain, blocking mitochondrial respiration.

**Figure 3.**
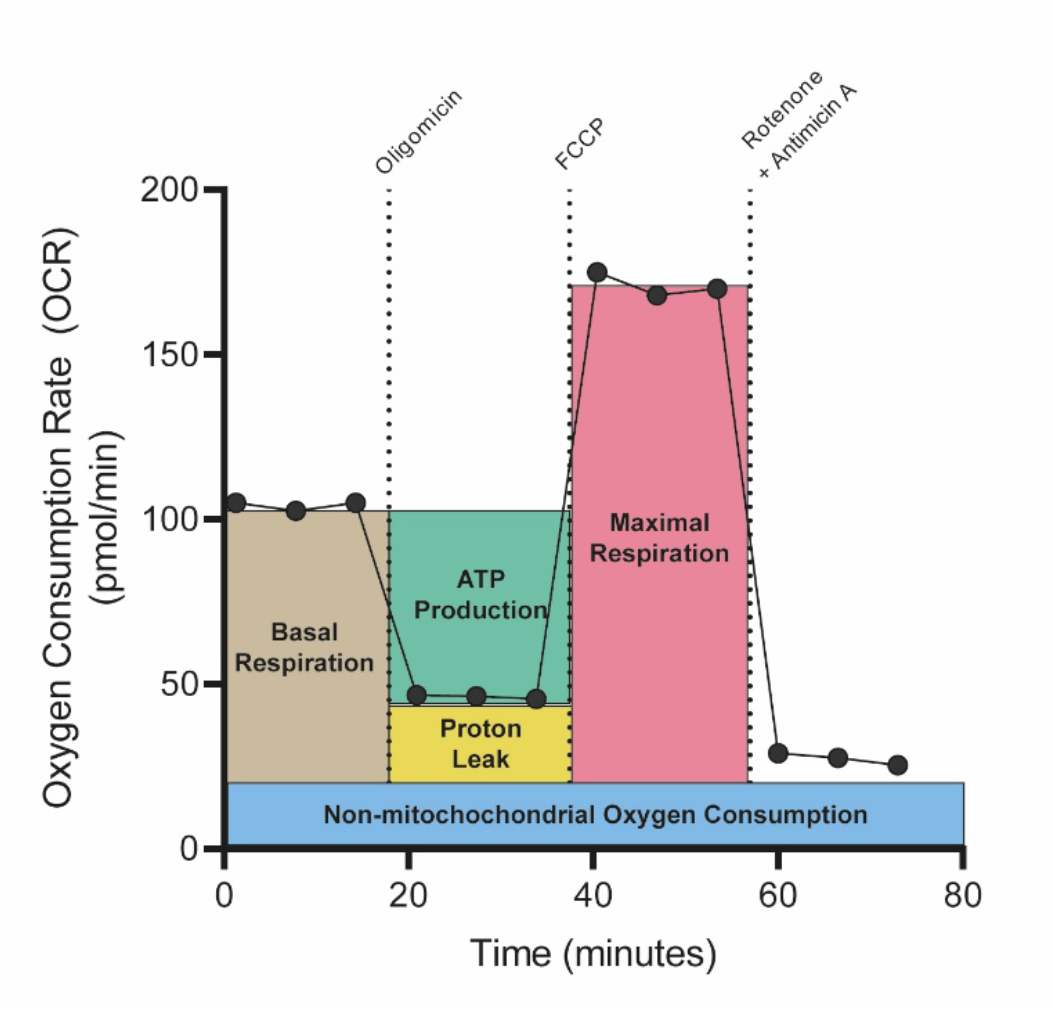
Seahorse Cell Mito Stress Test. Graphic representation of the different mitochondrial respiration parameters measured with the Seahorse XF Cell Mito Stress Test. Parameters were calculated as described. Non-mitochondrial respiration: minimum rate measurement after Rotenone/Antimycin A injection; basal respiration: (last rate measurement before first injection) – (non-mitochondrial respiration); maximal respiration: (maximum rate measurement after FCCP injection) – (non-mitochondrial respiration); proton leak: (minimum rate measurement after Oligomycin injection) – (non-mitochondrial respiration); ATP production: (last rate measurement before Oligomycin injection) – (minimum rate measurement after Oligomycin injection). Adapted from Agilent Technologies’ commercial materials.

Astrocyte and microglial cell cultures from WT and *Sfrp1*^-/-^ mice, were analyzed in control conditions and after 24-hour exposure to LPS (0.1 µg/mL). In homeostatic conditions, there was no major difference in mitochondrial respiration between WT and *Sfrp1*^-/-^ astrocytes (Fig. 4A, C–F). However, in response to LPS, *Sfrp1*^-/-^ astrocytes presented a consistent, although not statistically significant, decline in basal and maximal respiration (Fig. 4C-D), proton leak (Fig. 4E), and ATP production (Fig. 4F), compared to their WT counterparts. ECAR values in *Sfrp1*^-/-^ astrocytes were also somewhat lower than those of WT cells *Sfrp1*^-/-^. This difference became much more evident and statistically significant when the two genotypes were compared after LPS treatment (Fig. 4B; WT + LPS vs *Sfrp1*^-/-^ + LPS, *p < 0.05; **p < 0.01; multiple unpaired t-test). This data indicates that SFRP1 facilitates metabolic reprogramming of activated astrocytes toward glycolysis, an essential shift that supports their energy needs under stress conditions.

**Figure 4.**
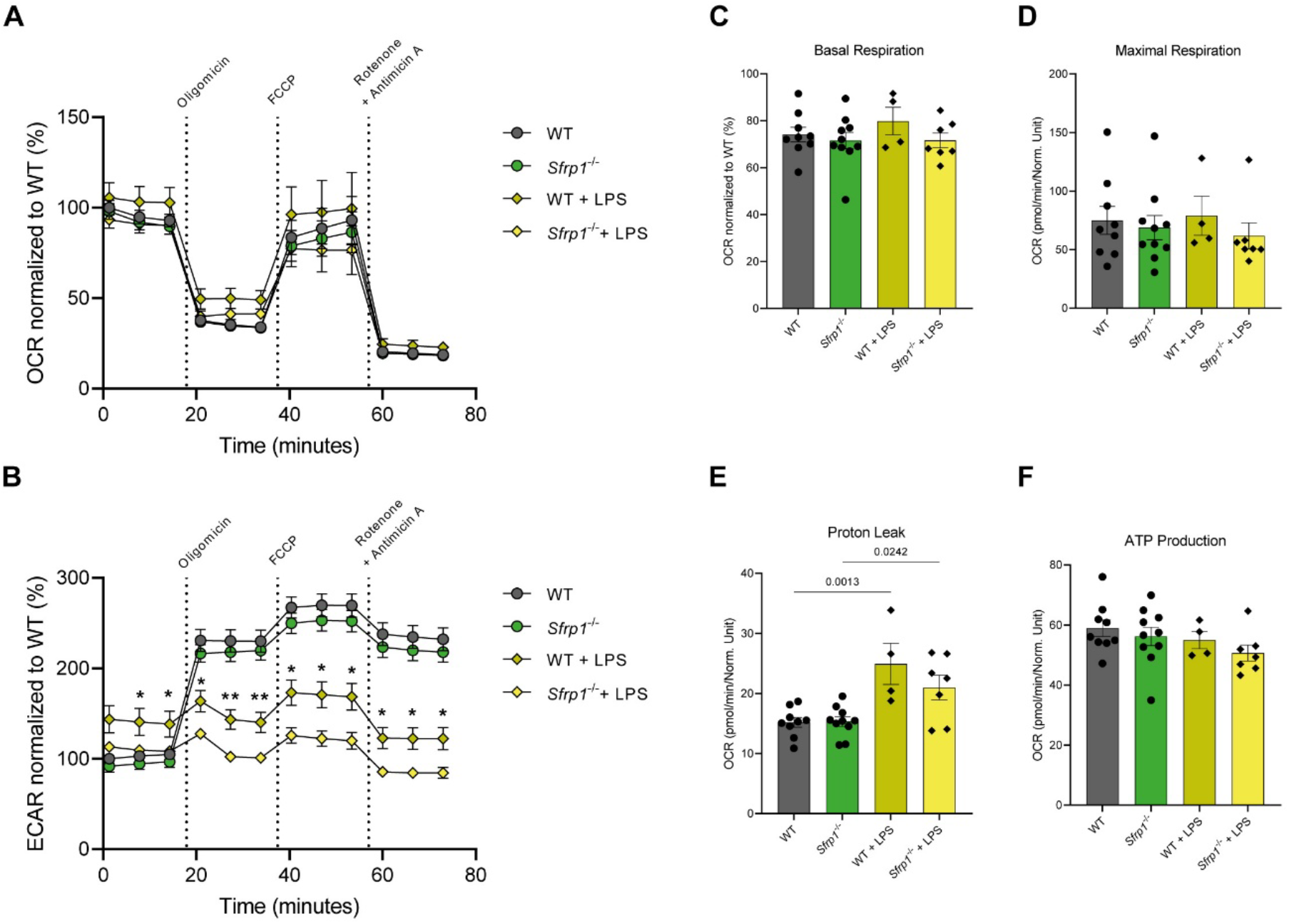
SFRP1 supports astrocytic glycolytic activity under LPS stimulation. The graphs show OCR (A) and ECAR (B) profiles measured by Seahorse analysis in astrocyte cultures, derived from WT and *Sfrp1*^-/-^ mice, treated with or without LPS (0.1 µg/mL) for 24 h. (C-F) Quantification of mitochondrial respiration parameters normalized to WT: (C) basal respiration, (D) maximal respiration, (E) proton leak and (F) ATP production. Values in the graphs correspond to the mean ± SEM of independent cultures (WT: n = 9; WT + LPS: n = 4; *Sfrp1*^-/-^ : n = 10; *Sfrp1*^-/-^ + LPS: n = 7), normalized to WT. Each measurement was performed using at least four technical replicates. Statistical comparisons were calculated with multiple unpaired t-test, followed by False Discovery Rate (FDR) correction (control for multiple comparisons) when compared WT + LPS vs *Sfrp1*^-/-^ + LPS, * p < 0.05; ** p < 0.01 for panels (A, B) and with one-way ANOVA followed by Šídák’s multiple comparisons test for panels (C-F) and p-values are indicated on the figure. ECAR: Extracellular Acidification Rate; LPS: Lipopolysaccharide; OCR: Oxygen consumption rate.

Microglial cells do not express *Sfrp1* themselves, but SFRP1 contributes to their response to an inflammatory stimulus *in vivo* or when co-cultured with astrocytes (Rueda-Carrasco et al., 2021). Consistent with this notion, it was perhaps not surprising to observe no meaningful difference in either mitochondrial respiration or glycolysis in isolated microglial cell cultures from *Sfrp1*^-/-^ and WT mice (Fig. 5A, B). While OCR values in knockout cells were slightly lower and more variable (Figure 5C, E, F), none of the changes were statistically significant. Similarly, LPS triggered a strong glycolytic response in microglial cells, consistent with already reported shift upon similar stimulation (Sabogal-Guáqueta et al., 2023), but there were minimal differences between the two genotypes.

**Figure 5.**
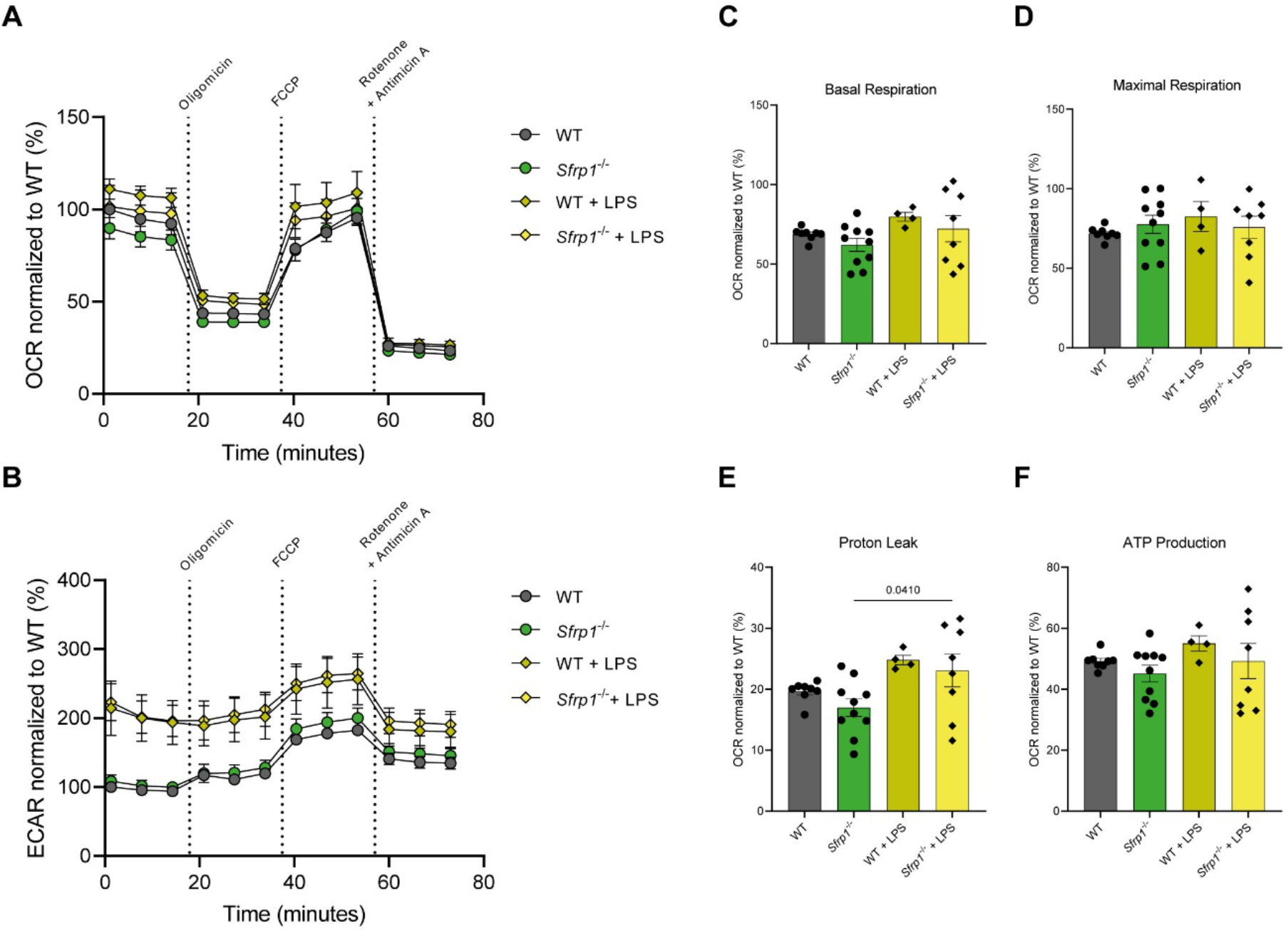
SFRP1 is dispensable for microglial mitochondrial and glycolytic functions. OCR (A) and ECAR (B) profiles measured by Seahorse analysis in microglial cell cultures, derived from WT and *Sfrp1*^-/-^ mice, treated with or without of LPS 0.1 (µg/mL) for 24 h. (C-F) Quantification of mitochondrial respiration parameters normalized to WT: (C) basal respiration, (D) maximal respiration, (E) proton leak and (F) ATP production. Values in the graphs correspond to the mean ± SEM of different independent cultures (WT: n = 8; WT + LPS: n = 4; *Sfrp1*^-/-^: n = 10; *Sfrp1*^-/-^ + LPS: n = 8), normalized to WT. Each measurement was performed using at least four technical replicates. Statistical comparisons were calculated with multiple unpaired t-test, followed by False Discovery Rate (FDR) correction (control for multiple comparisons) when compared WT + LPS vs *Sfrp1*^-/-^ + LPS for panels (A, B) and with one-way ANOVA followed by Šídák’s multiple comparisons test for panels (C-F) and p-values are indicated on the figure. ECAR: Extracellular Acidification Rate; LPS: Lipopolysaccharide; OCR: Oxygen consumption rate.

Altogether, these findings suggest that SFRP1 participates in the metabolic response of astrocytes to inflammation, pushing them towards glucose consumption.

### Increased SFRP1 levels boost glycolytic reservoir consumption in astrocytes

Astrocytes secrete SFRP1 into the culture medium and this secretion is significantly increased in the presence of microglial cells, which do not express but respond to its presence (Rueda-Carrasco et al., 2021). We thus explored whether addition of exogenous SFRP1 is sufficient to change astrocytes and microglial metabolism in the absence of an inflammatory stimulus, by treating both WT and *Sfrp1*^-/-^ astrocytes or microglia with hrSFRP1 (500 ng/mL) for 24 hr.

hrSFRP1 did not significantly modify mitochondrial respiration in WT and *Sfrp1*^-/-^ astrocytes, with OCR values and other oxidative phosphorylation parameters falling very closely to one another across all tested conditions (Fig. 6A, C–F). ECAR values, instead, rose in the presence of hrSFRP1, only mildly in WT astrocytes, but with significant, and timely sustained differences in *Sfrp1*^-/-^ cells (Fig. 6B; # p < 0.05, multiple unpaired t-test comparing *Sfrp1*^-/-^ vs *Sfrp1*^-/-^ + hrSFRP1). This effect became even more evident after oligomycin treatment, which blocks ATP synthase and forces cells to rely more heavily on glycolysis (Fig. 6B), further supporting the idea that SFRP1 fosters a glycolytic metabolism in astrocytes.

**Figure 6.**
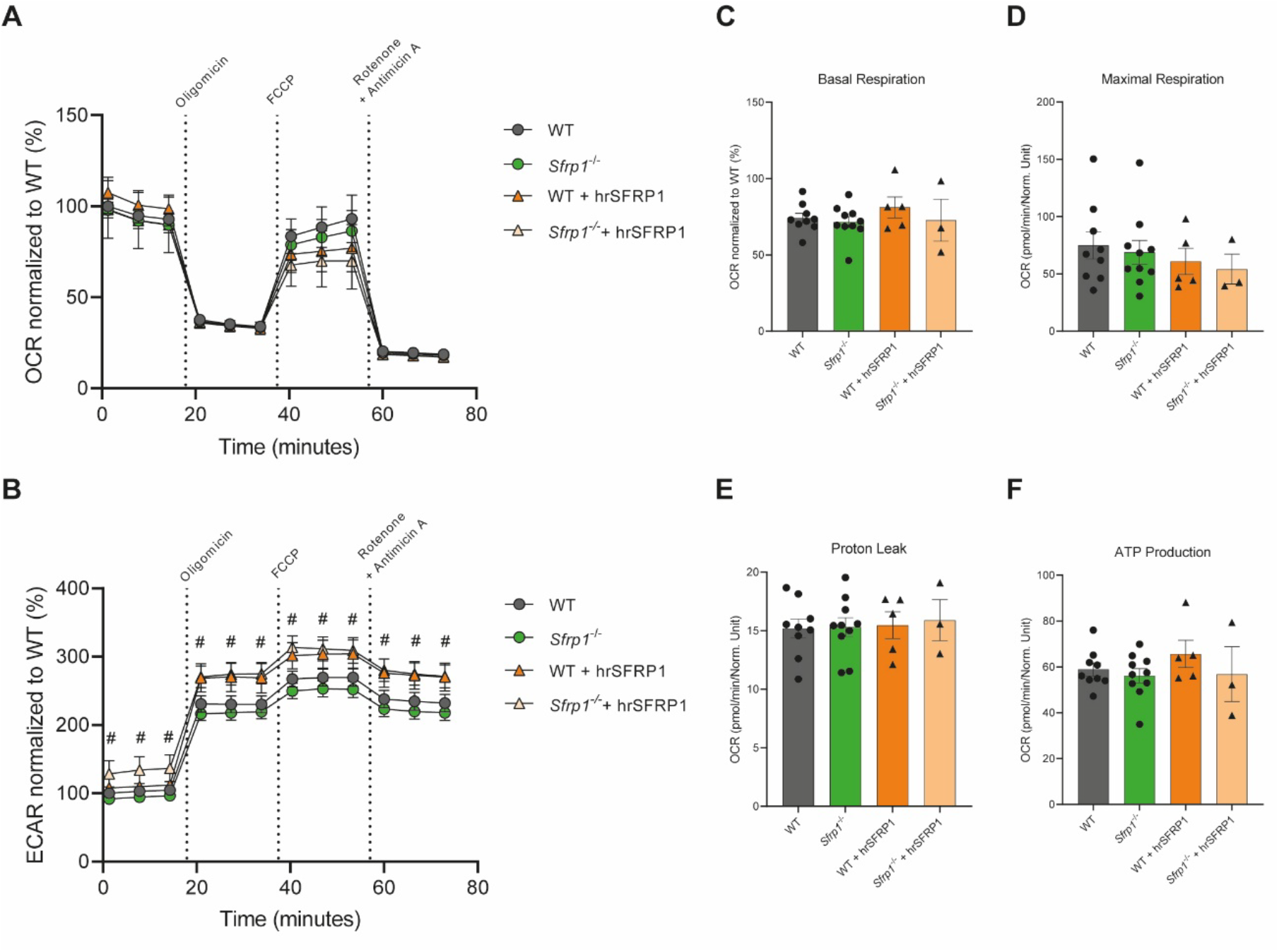
SFRP1 is sufficient to promote glycolysis in astrocytes. OCR (A) and (B) ECAR (B) profiles measured by Seahorse analysis in astrocytic cell cultures, derived from WT and *Sfrp1*^-/-^ mice, treated with or without hrSFRP1 (500 ng/mL) for 24 h. (C-F) Quantification of mitochondrial respiration parameters normalized to WT: (C) basal respiration, (D) maximal respiration, (E) proton leak and (F) ATP production. Values in graphs correspond to the mean ± SEM of independent cultures (WT: n = 9; WT + hrSFRP1: n = 5; *Sfrp1*^-/-^: n = 10; *Sfrp1*^-/-^ + hrSFRP1: n = 3), normalized to WT. Each measurement was performed using at least four technical replicates. Statistical comparisons were calculated with multiple unpaired t-test, followed by False Discovery Rate (FDR) correction (control for multiple comparisons) when compared *Sfrp1*^-/-^ vs *Sfrp1*^-/-^ + hrSFRP1, # p < 0.05, for panels (A, B) and with one-way ANOVA followed by Šídák’s multiple comparisons test for panels (C-F). ECAR: Extracellular Acidification Rate; OCR: Oxygen consumption rate.

By contrast, similar experiments in microglial cells revealed only minimal metabolic changes, although the ECAR values were quite variable among the different experiments (Fig. 7), making their interpretation difficult. *Sfrp1*^-/-^ microglia showed slightly reduced basal respiration, proton leak, and ATP production (Fig. C, E, F), while maximal respiration appeared to increase moderately (Fig. 7D). These subtle shifts might reflect a lower energy demand or a less activated state, at least in tested conditions that lack the *in vivo* interplay with astrocytes and other cell types.

**Figure 7.**
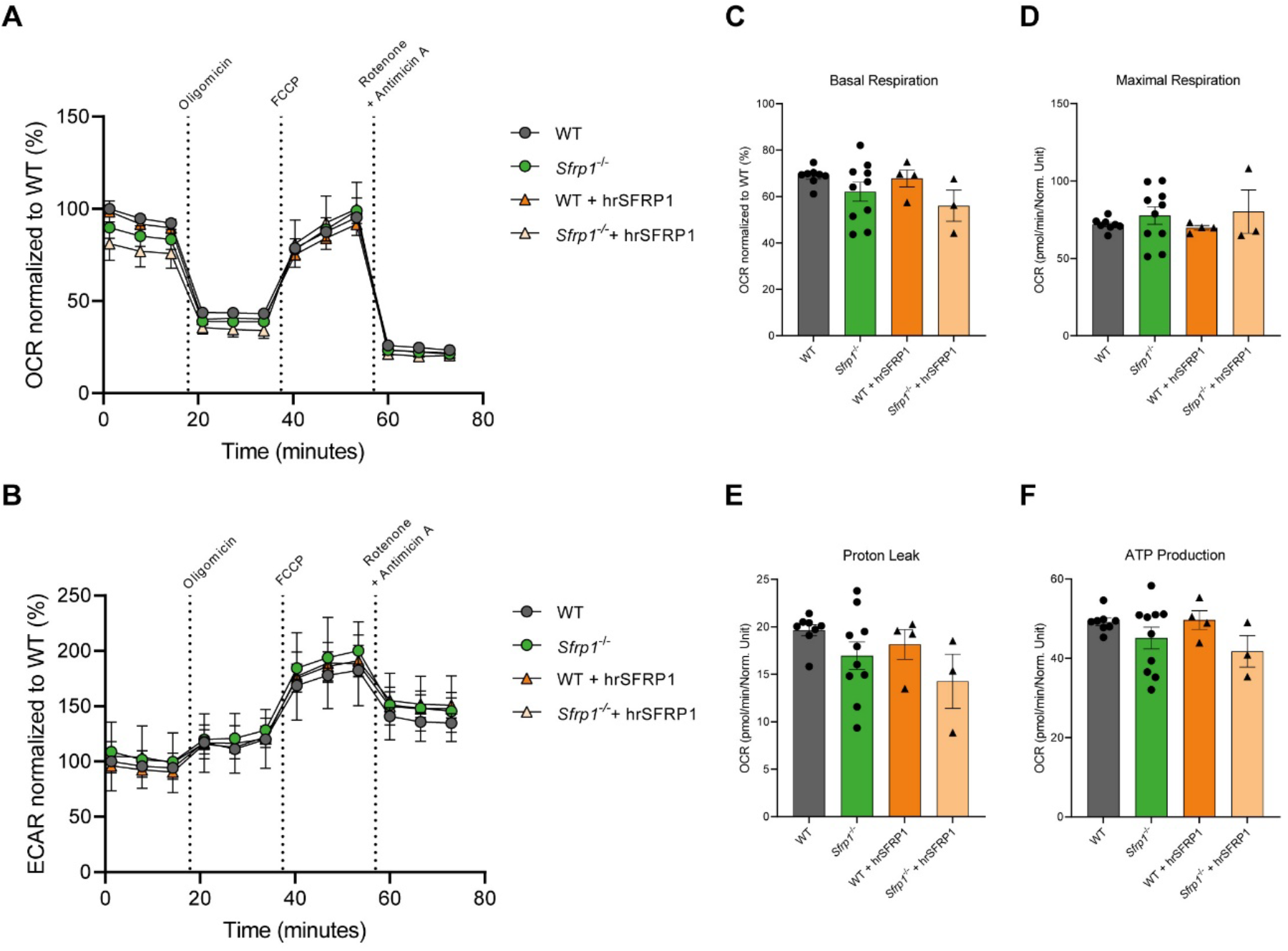
SFRP1 has no activity on metabolic profile of microglial cells. OCR (A) and ECAR (B) profiles measured by Seahorse analysis in microglial cell cultures, derived from WT and *Sfrp1*^-/-^ mice, treated with or without hrSFRP1 (500 ng/mL) for 24 h. (C-F) Quantification of mitochondrial respiration parameters normalized to WT: (C) basal respiration, (D) maximal respiration, (E) proton leak and (F) ATP production. Values in graphs correspond to the mean ± SEM of different independent cultures (WT: n = 9; WT + hrSFRP1: n = 4; *Sfrp1*^-/-^: n = 10; *Sfrp1*^-/-^ + hrSFRP1: n = 3), normalized to WT. Each measurement was performed using at least four technical replicates. Statistical comparisons were calculated with multiple unpaired t-test, followed by False Discovery Rate (FDR) correction (control for multiple comparisons) for panels (A, B) and with one-way ANOVA followed by Šídák’s multiple comparisons test for panels (C-F). ECAR: Extracellular Acidification Rate; OCR: Oxygen consumption rate.

### SFRP1 is sufficient to promote an energetic metabolism in astrocytes

To generate a comprehensive view of the energy shifts the two cell types undergo, and to compare their response to LPS or SFRP1 exposure, we plotted the obtained baseline OCR and ECAR values onto an “energy map”. This map allows the categorization of the metabolic state of the cells into four groups: glycolytic, quiescent, aerobic, and energetic (Ferrick et al., 2008). Under resting conditions, both astrocytes and microglia from the mutant mice clustered near their WT counterparts (Fig 8A), though *Sfrp1*^-/-^ astrocytes leaned slightly more towards a quiescent state (Fig 8A, C), while *Sfrp1*^-/-^ microglia slightly drifted toward a glycolytic state (Fig. 8B, D). Upon LPS stimulation, WT astrocytes shifted their metabolism towards an energetic state, marked by a clear rise of ECAR values and a more modest increase of OCR. By contrast, *Sfrp1*^-/-^ astrocytes rose their ECAR value only modestly and this was associated with a decrease in their OCR, so that their overall metabolic state is glycolytic rather than energetic (Fig. 8A). SFRP1 addition had a somewhat similar effect on WT astrocytes, promoting a more energetic state, although with a more balanced increase of both mitochondrial respiration and glycolytic consumption (Fig. 8C). *Sfrp1*^-/-^ astrocytes presented instead a more glycolytic profile (Fig. 8C), as observed with LPS addition.

**Figure 8.**
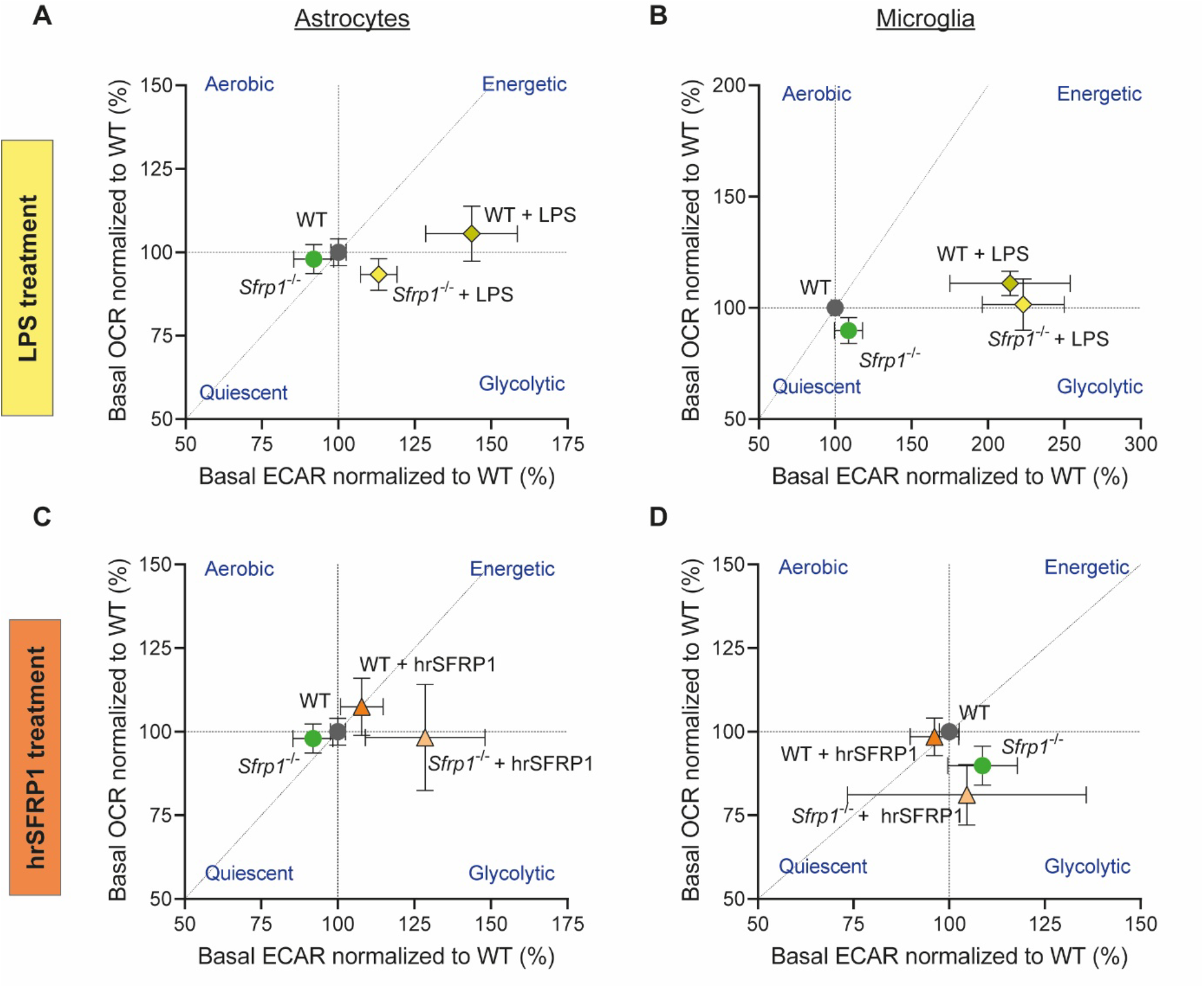
SFRP1 boosts astrocytes but not microglial cells toward an energetic profile. Representative energy map of astrocytes (A, C) and microglial cells (B, D), derived from WT and *Sfrp1*^-/-^ mice, treated with or without LPS (0.1 µg/mL; A, B) or hrSFRP1 (500 µg/mL; C, D) for 24 h. Values in all graphs correspond to the basal ECAR and OCR (mean ± SEM) calculated from the three time points, averaged across four technical replicates from independent cultures (WT: n = 8; WT + LPS: n = 4; *Sfrp1*^-/-^: n = 10; *Sfrp1*^-/-^ + LPS: n = 8; WT: n = 9; WT + hrSFRP1: n = 4; *Sfrp1*^-/-^: n = 10; *Sfrp1*^-/-^ + hrSFRP1: n = 3). Data were normalized to WT group. ECAR: Extracellular Acidification Rate; LPS: Lipopolysaccharide; OCR: Oxygen consumption rate.

The metabolic state of microglia from the mutant mice was very close to that of the WT, although with a slight tendency toward a glycolytic state, which was further promoted in both WT and mutant cells after LPS (Fig. 8B) or SFRP1 (Fig. 8D) stimulation.

Taking these finding altogether, we propose that SFRP1 is sufficient to promote a more energetic state in astrocytes, mostly influencing glycolytic activity rather than mitochondrial respiration. This effect is particularly evident when oxidative phosphorylation is limited (increased ECAR response after oligomycin exposure), thereby enhancing metabolic flexibility when it is most needed. By contrast, SFRP1 alone seems to have little influence on microglia metabolic response, suggesting that other astrocyte-derived factors, might predispose microglial response to SFRP1 *in vivo*, as we previously reported (Rueda-Carrasco et al., 2021).

## Discussion

In response to inflammation or injury, astrocytes and microglial cells undergo profound metabolic reprogramming, which contributes to restoring homeostasis and supporting neuronal function (Afridi et al., 2022). These metabolic changes must be integrated with other glial responses to damage, such as cytokine release or phagocytosis, implying the existence of specific mediators that coordinate astrocyte–microglia crosstalk, including their metabolic adaptations. In this context, we investigated whether SFRP1 can modulate glial cells metabolic adaptations, since previous studies, including ours, have shown that SFRP1 plays an important role in regulating inflammatory processes in different organs (Foronjy et al., 2010; Matsuyama et al., 2014; Rai et al., 2022; Yao et al., 2024), including the brain (Esteve, Rueda-Carrasco, et al., 2019; Rueda-Carrasco et al., 2021). Notably, SFRP1 influences the expression of a number of HIF and NF-κB target genes, many of which are directly linked to metabolic pathways. Our results demonstrate that SFRP1 acts as a driver of astrocytic metabolic activation, preferentially enhancing glycolysis over mitochondrial respiration. This effect is particularly evident under conditions of limited oxidative phosphorylation (inflammatory conditions), where SFRP1 boosts glycolytic flexibility to sustain energy production. By contrast, and to our surprise, SFRP1 alone has little impact on microglial metabolism, suggesting that additional signals, likely astrocyte-derived, are required to prime microglia for SFRP1 responsiveness, as we have previously reported *in vivo* (Rueda-Carrasco et al., 2021).

This conclusion is based the observation that *Sfrp1*-deficient astrocytes fail to upregulate their glycolytic activity in response to LPS, although it has little influence on glycolytic flux or mitochondrial respiration in homeostatic conditions. Furthermore, addition of human recombinant SFRP1 to cultured WT and *Sfrp1*^*-/-*^ astrocytes is sufficient to promote a higher glycolytic activity especially in *Sfrp1*^*-/-*^ astrocytes, further suggesting that SFRP1 normally acts through autocrine/paracrine signaling, influencing not only its cellular source, the astrocyte, but also cells present in the surrounding environment, such as microglial cells. Indeed, our study was motivated by the evidence that SFRP1 had an impact on microglia HIF1α and NF-κB signaling (Rueda-Carrasco et al., 2021). Unexpectedly, SFRP1 exerted only a subtle effect on microglial mitochondrial respiration. Microglia chronically deprived of SFRP1 (isolated from *Sfrp1*^-/-^ mice) exhibited a slight reduction in OCR under both basal conditions and following LPS stimulation, while glycolytic activity remained largely unaffected in either context. In both humans and mice, microglia undergo marked metabolic reprogramming upon LPS stimulation, characterized by a conserved upregulation of glycolytic genes such as HK1 and PFKFB3 downstream of TLR4 activation (Sabogal-Guáqueta et al., 2023). In mice, this reprogramming involves a shift from oxidative phosphorylation to a predominantly glycolytic phenotype (Geric et al., 2019; Orihuela et al., 2016; Suzuki et al., 2021), which, in turn, enhances NF-κB signaling and cytokine production (Cheng et al., 2021). These observations support the notion that metabolic reprogramming constitutes a hallmark of innate immune activation in microglia.

The mild changes we have observed align with the reported metabolic shift of LPS-stimulated microglia towards glycolysis. However, in contrast to previous reports, we observed a simultaneous small OCR increase. This discrepancy may reflect context-dependent metabolic flexibility, differences in experimental set-up or even, and more likely, the lack of an essential and continuous astrocytes to microglia crosstalk, which cannot be recreated in the Seahorse apparatus. This experimental limitation may also explain the lack of clear changes on microglia metabolism upon SFRP1 addition, despite previous studies (Rueda-Carrasco et al., 2021).

Taken together, these observations allow us to speculate that additional components of the astrocytes-microglia crosstalk are crucial for driving microglia metabolic reprogramming, and that such priming may be required to predispose microglial cell to respond effectively to SFRP1. Supporting this idea, Schwann cell-derived SFRP1 has been shown to regulate metabolic reprogramming and cytokine production in macrophages infiltrating injured peripheral nerves. Mechanistically, SFRP1 binds to both isoforms (α and β) of the macrophage Heat Shock Protein 90 (HSP90) (Yao et al., 2024), a chaperone known to modulate HIF1α and NF-κB in other contexts (Nagaraju et al., 2015). Notably, the HSP90α isoform is stress-inducible and itself regulated by NF-κB (Ammirante et al., 2008), raising the possibility that a similar feed-back loop may operate in microglia, enabling SFRP1-dependent metabolic responses only under specific inflammatory conditions.

Astrocyte and microglia metabolic reprogramming is often accompanied by mitochondrial structural remodeling. In hiPSC-derived astrocytes, chronic exposure to the neurotoxic amyloid-β peptide triggers neuroinflammation, shifts metabolism toward glycolysis fatty acid β-oxidation, and promotes a substantial mitochondrial remodeling with a transition from fusion to excessive fission and swelling (Zyśk et al., 2023). Similarly, LPS stimulation leads to mitochondrial fragmentation in astrocytes, associated with increased DRP1 and autophagic processes activation, essential for preserving mitochondrial network integrity (Motori et al., 2013). In contrast, LPS-mediated TLR4 activation in microglia promoted initial fission followed by Mfn2 and DRP1 mediated compensatory fusion (Katoh et al., 2017), which maintains metabolic efficiency during stress (Adebayo et al., 2021). In our study, hrSFRP1 treatment enhanced glycolytic activity without altering mitochondrial number or network morphology in astrocytes, whereas SFRP1-deficient microglia exhibited fewer individual mitochondria and elongated network branches. Notably, however, hrSFRP1 treatment did not affect microglial mitochondrial distribution, suggesting that these structural features may reflect developmental or differentiation-related adaptations, potentially linked to the absence of SFRP1-mediated inhibition of Wnt/β-catenin signaling, which is known to favor mitochondrial fusion over fission (Brown et al., 2017). This finding, consistent with OCR Seahorse measurements, suggests that SFRP1 alone is not sufficient to drive mitochondrial remodeling in astrocytes, at least in the absence of an inflammatory stimulus.

In conclusion, SFRP1 appears to induce robust changes in astrocytes glycolytic activity, while other parameters were only slightly modified after LPS stimulation with trends that, however, are largely consistent with previous published data. This alignment likely lends biological relevance to our finding, especially considering the limitations of our experimental set-up. Indeed, primary cultures of astrocytes and microglia are inherently variable, both between preparations and across experimental replicates, introducing noise that can mask subtle but meaningful effects. The Seahorse assay is known to exhibit technical variability itself (Mercier-Letondal et al., 2021; Yépez et al., 2018), especially in primary cultures (Hammoud et al., 2025), and this further complicates the interpretation of mitochondrial respiration data. Morphological analysis of mitochondria is also subject to considerable heterogeneity, both at the cellular and mitochondrial levels, posing significant challenges in obtaining statistically strong conclusions. Future studies with higher experimental power and complementary techniques will be necessary to confirm the trends observed in our analysis. Nevertheless, our study unveils a previously unknown function of SFRP1 on astrocytic metabolic reprogramming during neuroinflammation. A deeper understanding of this pathway may provide new perspectives on how astrocyte metabolism is tuned in the inflamed brain and further support the idea that SFRP1 might be multifaced target for complex neurodegenerative conditions, such as Alzheimer’s disease, in which SFRP1 seems to play multiple roles (Esteve, Rueda-Carrasco, et al., 2019; Pereyra et al., 2025).

## Author Contributions

MMB, GP and PB set up the research project and designed the experiments. MMB and MJMB conducted experiments and data acquisition. PM contributed to *in vitro* experiments. MMB analyzed the data. MMB and PB wrote the manuscript. PB secured funding and supervised the work.

## Acknowledgments

We thank Andrés Vicente Acosta for assisting with Seahorse technique and analysis, and members of the CBM SMOA and Cytometry facilities for their support and advice.

